# Temporal Processing during Decision Making under Uncertainty

**DOI:** 10.64898/2026.06.30.735611

**Authors:** Rikki Rabinovich, Daniel Feldman, Rhiannon Cowan, T. Alexander Price, Niloufar Shahdoust, Tyler S. Davis, Shervin Rahimpour, Ben Shofty, Elliot H. Smith

## Abstract

Time perception and decision-making are interrelated processes that are critical to much of human and animal behavior. Although the behavioral importance of this link between the processes has been established, the neural underpinnings of temporal decision-making are unclear. The current study leverages human intracranial electrophysiology to investigate the neural contribution of temporal processing to decision-making under risk. To probe temporal decision-making, we use a version of the Balloon Analogue Risk Task, in which participants inflate virtual balloons with the goal of reaching maximal balloon size but stopping inflation before the balloon pops. Points are awarded based on inflation duration of successful trials, such that points awarded are linearly related to balloon size. By decoding temporal features of the task on a trial-by-trial basis, we find neural representations of time during balloon inflation. Specifically, we find that both medial and lateral temporal regions of the brain encode the majority of this time-related signal. Importantly, we can decode the progression of time leading up to the outcome of a successful decision (pressing a button to stop inflation before the balloon pops). This encoding of time depends on an active decision to end the trial: we cannot decode time during passive observation of balloon inflation to a known maximal size. To validate these results further, we use a temporal convolution network to predict participants’ response (i.e., time of button press) based on the brain activity leading up to the response. We find that we can predict the moment of response with high accuracy and temporal resolution. The model predicts the timing of successful decisions but not the timing of balloon pops. Together, these findings reveal neural representations that encode the progression of time, specifically in relation to an active decision.

## Introduction

Integrating temporal and sensory information is essential to build and update models of the external environment and internal state (Clark, 2013), making timing a vital component of predictive decision-making. The importance of time-keeping has led to a long-time search for neural “clocks” or other neural mechanisms for temporal processing. Various candidate regions have been implicated—from cerebellum (Ashe and Bushara, 2014) to medial temporal regions (Pastalkova et al., 2008; Tsao et al., 2018; Umbach et al., 2020) to striatum (Bakhurin et al., 2017; Mello et al., 2015; Toso et al., 2021), and even sensory cortex (Rabinovich et al., 2022; Salvioni et al., 2013). However, these studies focus mainly on neural correlates of externally imposed time intervals, while ethological decision-making often involves self-generated durations, such as deciding *when* to execute an action.

Indeed, time and decision-making are inextricably linked, with both processes bidirectionally influencing each other. For instance, decision-making under time pressure has been shown to enhance risk-seeking behavior (Young et al., 2012), while conversely, impulsive traits influence an individual’s perception of time (Wittmann and Paulus, 2008). Although existing literature describes this behavioral relationship between time perception and decision-making, the neural basis of human temporal decision-making is still unknown—partly due to limitations of conventional fMRI approaches, which lack the temporal resolution needed to track rapidly unfolding neural dynamics that subserve precisely timed decisions.

The present study sought to uncover temporal processing in the human brain during risk evaluation and timing-dependent decision making. To this end, we leveraged high temporal resolution intracranial electrophysiology in the human brain, and applied a version of the Balloon Analogue Risk Task (BART). Traditionally used to study impulsive behavior and risk-seeking (Lejuez et al., 2002; Rao et al., 2008), this task is also ideal for investigating temporal decision-making. Unlike other decision-making tasks, for which accuracy is the primary measure and timing is secondary, in BART, high task performance requires accurate time interval estimation and successful learning of the relationship between time interval and sensory stimulus.

Note that BART does not entirely isolate interval timing from sensory feedback: a visual stimulus that evolves at a constant rate enables the participant to predict *when* to act during each trial. Rather than eliminating the need for time estimation, visual feedback helps reorient the participant, anchoring their time perception and improving temporal predictions.

From intracranial local field potential (LFP) recordings, we uncovered not only risk representations during BART performance, but also encoding of decision-related time information. Notably, neural activity encoded the linear passage of time only when it was behaviorally relevant and predicted the timing of planned, self-generated decisions, rather than passive or externally-driven trial events.

## Subjects

### Methods

18 neurosurgical patients undergoing clinical monitoring for epilepsy participated in the study. All subjects provided informed consent prior to participation, both for implantation of microelectrodes and for participation in cognitive tasks. All experimental procedures were approved by the Institutional Review Board. Only patients aged 18 and over participated in the experiments. No patients with major neuroanatomical anomalies or cognitive deficits were included.

### Behavioral Paradigm

Patients performed a version of the Balloon Analogue Risk Task (BART; Fig. 1a). For each trial, a colored circle “balloon” appeared on the computer monitor, and subjects were instructed to press a button on a video game controller to start inflating a balloon. Each balloon could inflate up to a predetermined size (jittered across a probabilistic distribution) before popping. However, prior to the pop, subjects could choose to press the button again at any point to stop inflation. When subjects successfully stopped inflation before the balloon “popped”, they were awarded points ($). These successful trials were called *bank* trials. The value of the reward was linearly related to the duration of inflation, and hence the size of the balloon at the end of the inflation. If subjects did not stop inflation in time, and the balloon popped, no points were awarded. These error trials were *pop* trials.

**Figure 1.**
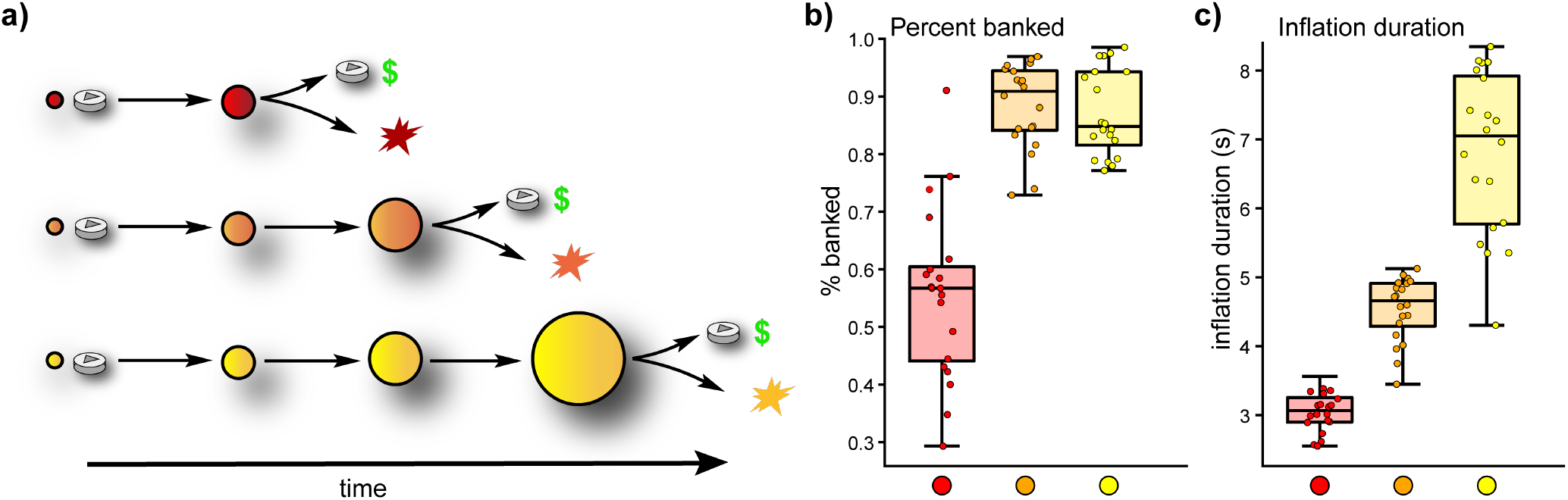
Experimental task and behavior. a) Schematic of the behavioral paradigm: subjects performed the Balloon Analogue Risk Task. For each trial, a red, orange, or yellow balloon appears on the screen (cue period). Then subjects press a button to begin inflation and choose when to press the same button again to stop inflation. If inflation is stopped before the balloon pops, subjects are awarded points ($). These successful trials are bank trials. The value of the reward corresponds to the size of the balloon at the end of the inflation. If subjects do not stop inflation in time, and the balloon pops, no points are awarded. Maximum balloon size differs across trials within a probabilistic range, based on balloon color (red: small; orange: medium; yellow: large). b) Behavioral performance quantified as the percent of successful (banked) trials for red, orange, and yellow balloons. Each dot is average data for that trial type for 1 session (N = 20 sessions). c) Behavioral performance quantified as the duration of inflation for each balloon color (only for bank trials).

Red balloons were programmed to pop at the smallest size, orange balloons popped at a medium size, and yellow balloons inflated the longest before popping. In addition to these active trials, there were also risk-free control trials, which were guaranteed to result in a bank outcome with no action required from the participant. During these trials, colored balloons appeared in the center of a grey ring; after the first button press, the balloons would inflate until they reached the diameter of the grey ring, at which point the trial was complete and points were awarded. There were approximately 3 times as many active trials as passive trials.

Each session had an average of 226 trials (range: 69 – 256). Reaction time (RT) was defined as the interval between balloon onset and the first button press initiating inflation. Trials with RT *>*5 seconds were excluded from CEBRA and SVM decoding, because they indicated disengagement from the task. Most commonly, these were the first few trials of the session, while patients were still learning the task structure.

### Electrophysiological recordings

Depth electrodes were implanted in patients’ brains as part of their epilepsy monitoring, and the location of the implants was dictated by clinical need. 0.8 mm diameter DIXI stereoelectroencephalography (sEEG) electrodes were used, containing macrocontacts with 3.5 mm spacing between contacts and 2 mm contact length. LFP recordings were conducted using a Blackrock Microsystems recording system (128 channels, 1 KHz sampling rate). 2 of the 18 patients performed 2 sessions on different days, so 20 sessions in total were collected. Recording sites were classified as belonging to a particular region and brain network using the LeGui software package (Davis et al., 2021), based on a post-operative CT scan and co-registered structural MRI and normalized to MNI space (warping from patient space to MNI space was required due to anatomical variations between subjects). Regions were identified using the Neuromorphometrics (NMM) atlas. Figure 2 illustrates the approximate locations of recording sites in various brain regions.

**Figure 2.**
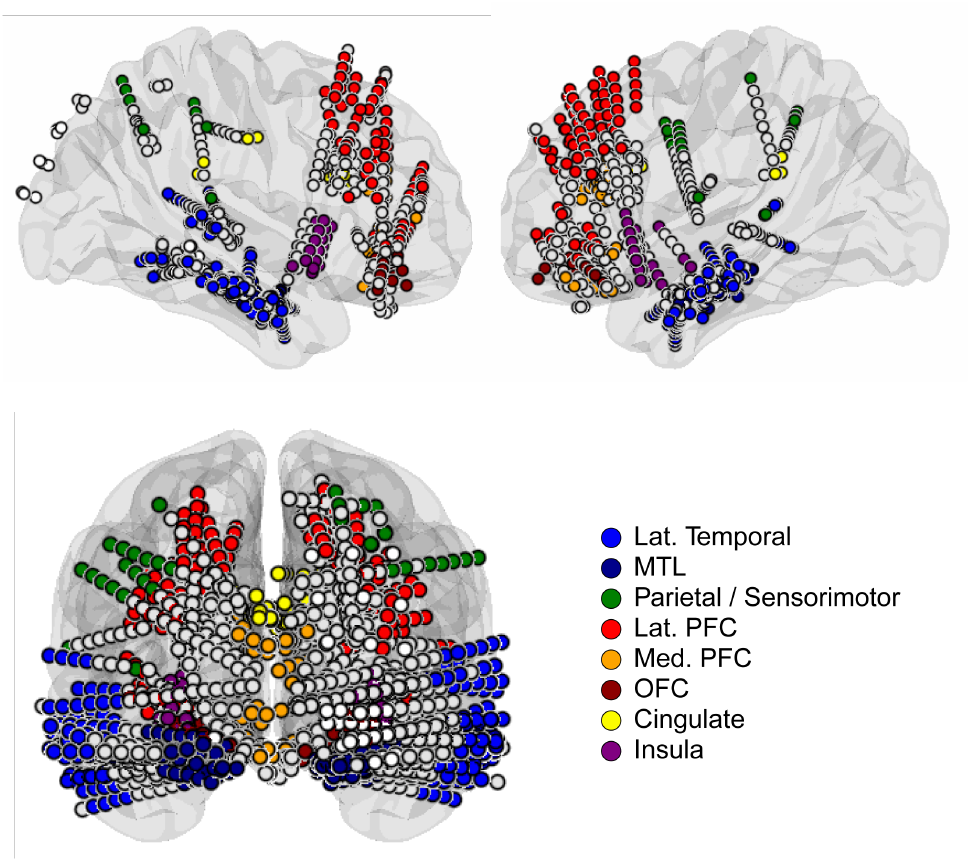
Signals are recorded from implanted sEEG electrodes during task execution. The model brain illustrates approximate recording locations of the full set of contacts for all patients (N = 18), warped from patient space to MNI space due to anatomical variations between subjects, and overlaid on an average transparent model brain . Linearly aligned contacts resided on the same electrode lead. Contacts are colored according to brain region. White contacts are either within white matter or in regions not discussed here. Lat. Temporal: Lateral Temopral; MTL: Medial Temporal Lobe; Lat./Med. PFC: Lateral/Medial Prefrontal Cortex; OFC: Orbitofrontal Cortex.

### Analysis of LFP power

The Python MNE library was used to analyze LFP data (Gramfort et al., 2013; Larson et al., 2025). Data pre-processing included removing noisy channels, notch filtering to remove power line noise (60 Hz and 120 Hz harmonic), and common average referencing. Data was epoched to only include relevant time windows. Frequency bands were split as follows: alpha (8 – 13 Hz), beta (13 – 30 Hz), gamma (30 – 70 Hz), and high gamma (70 – 140 Hz). Power spectra were calculated using the Welch method, with a window of 0.1 seconds.

### Decoding

#### Linear classification

We used Python scikit-learn software (Pedregosa et al., 2011) to train linear models to perform support vector classification to classify outcome or time bins. For outcome decoding, we subsampled to ensure equal numbers of bank and pop trials. When training and testing on different data (e.g., training on time bins around outcome; testing on earlier time bins), no cross-validation was performed. Otherwise, K-fold cross validation was performed within-subject for linear decoders: each dataset was divided into 4 test-train splits based on trial index; timepoints within the same trial were kept together. Linear classifiers were trained on a subset (3/4) of trials for each subject, for each fold. Classifiers were then tested on the held out trials. Classification performance was quantified using one-sample two-sided *t*-tests comparing each subject’s decoding accuracy to chance. Each timepoint and frequency band constituted a single feature. For linear decoders, we identified feature importance through Haufe-transformed coefficients of the linear SVM, using the MNE python library to extract spatial patterns from the coefficients Haufe et al., 2014.

#### Nonlinear dimensionality reduction and decoding

For nonlinear analysis, we applied the “Consistent EmBeddings of High-dimensional Recordings using Auxiliary variables” (CEBRA) technique, a recently developed nonlinear encoder and dimensionality reduction technique (Schneider et al., 2023). This method uses contrastive learning to identify latent structure in high-dimensional neural time-series data, pulling together related data points and pushing apart unrelated ones. With behavioral labels to supervise the model’s learning, the model successfully identified low-dimensional structure related to aspects of the BART task from the high dimensional neural data. We fit independent CE-BRA models on the data for each patient, then used the trained model to transform the data into 5-dimensional embeddings. We then aligned the embeddings across all patients with Procrustes alignment. For decoding, we used 4-fold cross-validation to train the supervised CE-BRA models: we randomly split the data into 4 folds, preserving temporal structure (timepoints within the same trial were not separated; only whole trials were held out). Train/test splits for each subject were chosen prior to training the models. We then trained the CEBRA models for each patient, holding out each fold (test set) while training on the remaining data (training set). Models trained on the training set were used to generate embeddings for the test set for each cross-validation fold. We then used these test embeddings for decoding the behavioral information related to the held out trials: for each fold, we used a K-nearest neighbors (KNN) classifier to classify relative or absolute time. Since we aligned embeddings across patients and only ran the decoder once per validation fold, we simply used a one-sample *t*-test to compare the performance on all cross-validation folds to chance level. Traditional permutation tests with CEBRA models were not feasible, given the size of the data and the required computing power.

#### Temporal convolutional network

To predict the timing of an imminent decision, we used a nonlinear causal temporal convolutional network (TCN). To accommodate minor temporal variability and sampling jitter, supervision was defined as a temporal window centered on the true event time rather than a single time point. Specifically, model targets were set to 1 within ±1 time step of the event index (corresponding to approximately ±100 ms); all other time steps were labeled as 0. This formulation reframes event detection as a dense temporal classification problem, allowing the model to learn distributed predictive signals that arise prior to the decision rather than relying on an instantaneous marker. The model architecture was a causal temporal convolutional network.

For each trial, channel activity was first aggregated using a masked mean across channels, yielding a time × frequency-band (alpha, beta, gamma, high gamma) representation. The pooled signal was then passed through a 1×1 convolution to project frequency bands into a higher-dimensional latent space. The model then consisted of a stack of 8 causal temporal convolutional blocks with exponentially increasing dilation factors between TCN blocks. A final 1×1 convolution was applied, producing a scalar logit at each time point for prediction. Each temporal convolutional block operated on 64 latent feature channels. Blocks comprised two causal 1D convolutions (kernel size = 5) with shared dilation, followed by batch normalization, GELU activation, and dropout (p = 0.2). Residual skip connections linked the input and output of each block. Dilation increased exponentially across the 8 blocks (2^0^–2^7^), enabling multi-scale temporal integration while preserving strict causality. Causal padding ensured that predictions at time t depended only on past neural activity. Models were trained using a masked binary cross-entropy loss. Optimization was performed using AdamW with gradient clipping to stabilize training.

Model performance was assessed with 4-fold cross validation. Performance was quantified by mean absolute error (MAE) between the predicted event time (maximum posterior probability) and the true event time, as well as accuracy defined as the proportion of trials for which the predicted event fell within the time window of the true event. These metrics jointly capture temporal precision and trial-level accuracy.

## Results

To study temporal processing in association with decision making, we recorded intracranial local field potentials (LFP) from the brains of 18 neurosurgical patients (Fig. 2) performing a risky time-dependent decision-making task (BART; Figs. 1; S1). Participants “inflated” virtual balloons (by pressing a button to start and stop inflation), with the goal of inflating each balloon as much as possible before it popped (Fig. 1a). Each balloon color was associated with a different degree of risk, based on its likelihood of popping at various sizes: red balloons popped at the smallest size, orange balloons were intermediate, and yellow balloons could inflate to the largest size before popping (Fig. S1b). For each balloon color, the maximum size at the time of the pop was stochastic and variable across trials (S1c). Overall, participants showed satisfactory behavioral performance on the task, especially for orange and yellow balloons, which allowed longer average inflation times (Figs. 1b,c; S1). Notably, inflation times, particularly for yellow balloons, exhibited substantial variation (Figs. 1c; S1a).

### Risk is assessed during stimulus presentation

First, we considered the neural representations of the three trial types (red, orange, and yellow balloon colors) at different time windows in the trial. We used the nonlinear encoder and dimensionality reduction tool CEBRA (Schneider et al., 2023) to generate low-dimensional embeddings corresponding to the three balloon colors, with color as the supervisory signal (Fig. 3a; Figs. S2, S3). We examined both the cue period and the final three seconds of the inflation period. For both cue and inflation, we considered only successful trials, so both analyses included the same trials. We found that the brain states clustered such that the trial types were separated (Fig. 3a), and we observed two notable features of the clustering. First, despite similar performance (percent banked) for yellow and orange balloons (Fig. 1b), the most distinct cluster of neural data was during the time bins related to viewing the yellow balloons. Second, the clustering was stronger for the cue period than during inflation (Fig. 3b). This result suggests a neural representation of risk that evolves over time but is particularly strong while assessing the cue and before initiating the inflation.

**Figure 3.**
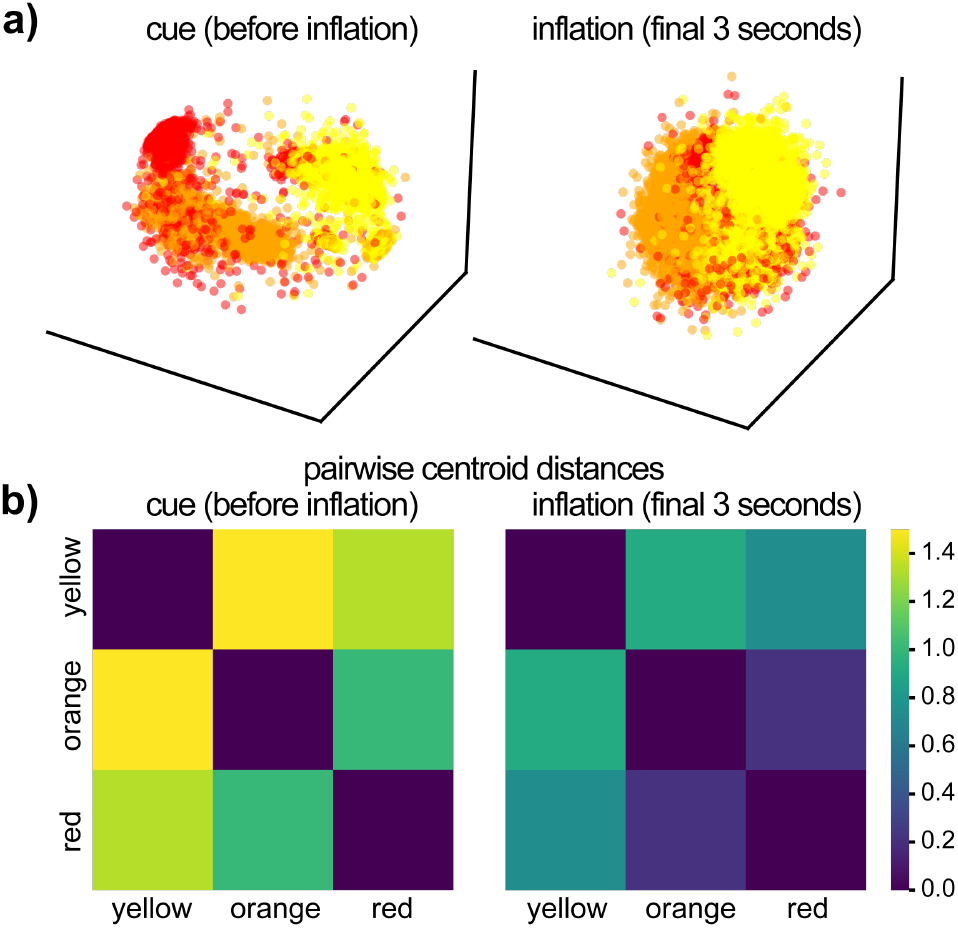
Neural representations of balloon colors during the cue period (pre-inflation, left) and during the final 3 seconds of inflation (right). a) Visualization of red, orange, and yellow trial types in a low-dimensional latent space generated by CEBRA models trained to distinguish balloon colors. b) Pairwise centroid distances between the clusters corresponding to the three balloon colors.

### Decision-related representations extend backwards in time

We investigated neural representations of trial-level outcome: *bank* (successful trials ended by the participant via a button press) or *pop* (unsuccessful trials that ended unexpectedly with a balloon pop before the subject’s decision to end the trial). We trained a linear support vector machine (SVM) to classify pop vs. bank trials (for active trials of all balloon colors) using neural activity from a 100 ms window centered on the outcome (50 ms prior – 50 ms after outcome). We then applied this trained classifier to non-overlapping data from the immediately preceding 100 ms window (150 – 50 ms before outcome). Within this temporally distinct window, outcome could be predicted with 66.1% accuracy (Fig. 4a; one-sample two-sided *t*-test, *p* = 2.13 × 10^−7^; bank trials: 62.67%, *p* = 1.17 × 10^−3^; pop trials: 69.08%, *p* = 1.07 × 10^−6^). Thus, outcome-related representations are already present in the moments leading up to the outcome, suggesting that these signals are related to the imminent temporal decision.

**Figure 4.**
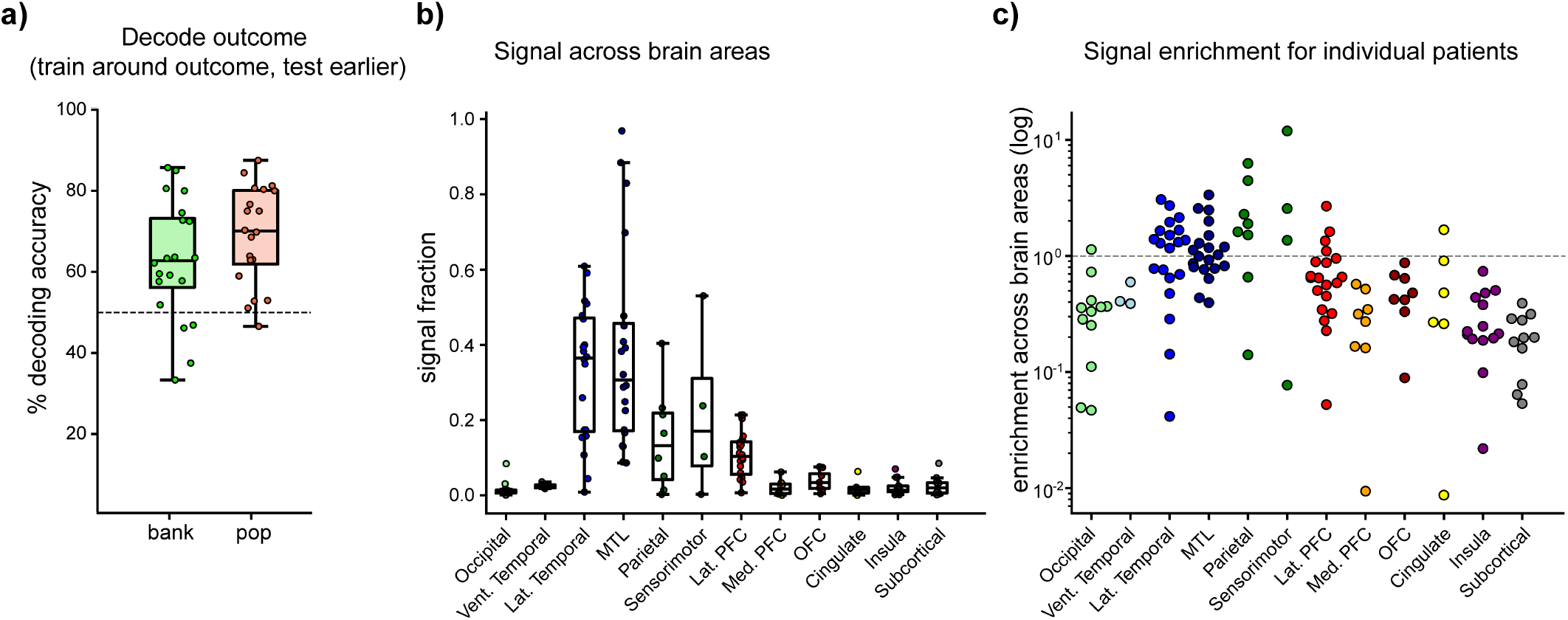
Decoding of outcome identity (N = 20 sessions) using a linear support vector classifier trained on bank vs. pop outcomes for each participant. a) Decoding performance for bank and pop trials. b) Signal carried by different brain regions, based on coefficients (spatial patterns) extracted from the trained model for each patient. Each dot is the average signal fraction (proportion of the total signal contained in each region) for a single session. c) Inter-patient variability in signal enrichment, or the degree to which the signal in each region is stronger than expected based on coverage density in that region. Each dot is one patient’s recording session.

Next, we sought to identify the neurophysiological correlates of this representation. We examined the activation patterns (Haufe-transformed coefficient weights of the linear classifier (Haufe et al., 2014)) to uncover the underlying spatial patterns that carry the decision-related information (Fig. 4b,c). We found the strongest signals in temporal regions—both lateral and medial temporal lobe (MTL). Parietal and sensorimotor areas also contributed to these anticipatory signals, albeit to a lesser extent, suggesting a sensorimotor component to the preparatory state. In sum, these findings not only reveal encoding of imminent decision, but also suggest that this decision-related signal is temporally-extended backward in time.

### Neuronal population activity tracks the progression of time

So far, we only considered aspects of the upcoming decision; next, we asked whether the progression of time was itself represented in the neural activity patterns. First, we examined the representations of relative time on bank trials, which started and ended with a button press. We used CEBRA to generate low-dimensional embeddings of the brain state trajectories over the course of the entire inflation period (Fig. 5a, left), with time-in-trial split into 10 bins (i.e., the initial button press to start inflation was labeled time 0, and the final button press to stop inflation was labeled time 9). To quantify the encoding of relative time, we trained CE-BRA models to predict time-in-trial, but used 4-fold trial-level cross-validation, see Methods and Supplemental Methods). After fitting a model on each training set, we performed the same transformation on the test set to generate a test embedding. From the test embeddings, we used a K-nearest neighbors (KNN) classifier to decode time-in-trial and saw above chance decoding performance (Fig. 5a, right; yellow balloons: one-sample two-sided *t*-test, *p* = 0.0031; red balloons: *p* = 0.0003), even for yellow-balloon trials, which had high variability in inflation times (Fig. S1a).

**Figure 5.**
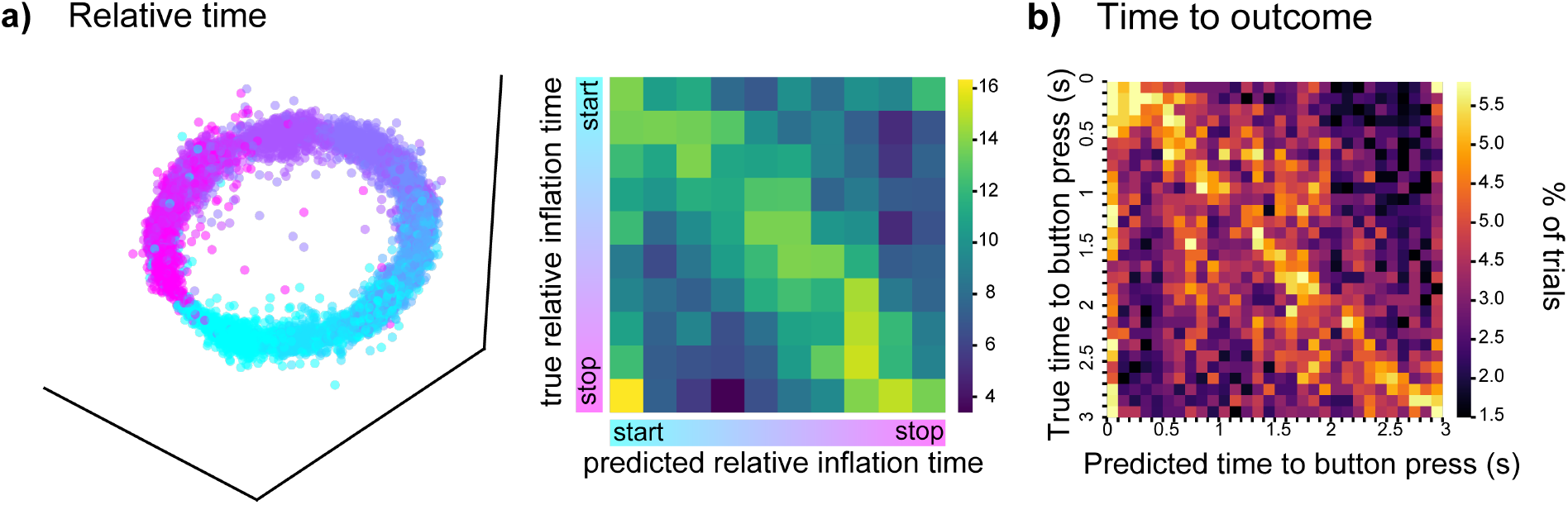
Decoding relative and absolute (outcome-aligned) time. a) CEBRA models were trained on neural signals over the course of the entire inflation duration for all banked yellow-balloon trials, with discretized time as the supervisory signal: each trial’s inflation period was split into 10 equal bins. Left: CEBRA-generated embeddings aligned across all participants from the entire neural dataset. Each dot corresponds to 1 time point (0.1 second bins). Color shows the spectrum from early to late inflation: blue dots indicate timepoints early during inflation, while pink dots are timepoints toward the end of inflation (closer to the final button press). Right: Decoding of relative time (discretized time bin) from embeddings generated from held-out data. The blue-yellow colorbar shows the percent of trials for which the time bin on the y-axis that were predicted to be the time bin on the x-axis. b) Decoding of absolute time-to-outcome based on cross-validated CEBRA embeddings. Models were trained to distinguish 0.1 second time bins during the 3 seconds of inflation leading up to the final button press of each banked yellow-balloon trial. Time 0 is the final time bin before the button press.

Since we found evidence of temporally-extended preparatory signals prior to a decision, as well as relative time representations during balloon inflation, we next asked if the neural activity also contained information about the absolute progression of time toward a decision. We trained CEBRA models on neural data during the 3 seconds leading up to the button press, with time-to-outcome (split into 0.1 second time bins) as the supervisory signal (Fig. S4). Here, we only analyzed yellow-balloon bank trials, which were long enough to ensure >3 second inflation for most trials but still displayed substantial variation in inflation time (Fig. S1a). Cross-validated model training followed by KNN decoding resulted in above-chance time decoding (Fig. 5b; *p* = 0.0015).

To determine whether this time-tracking relies on similar underlying neural processes as the encoding of the temporally-extended decision discussed above, we repeated the time-to-outcome decoding using a linear SVM for improved interpretability. A linear SVM could predict time-to-outcome with above-chance accuracy (two-sided one sample *t*-test, *p* = 0.00014 for 0.5 second time bins), corroborating the CEBRA results. Analysis of the spatial patterns underlying the time decoding (Fig. 6a) revealed neuroanatomical signal distribution similar to that of the temporally-extended outcome-identity decoder (Fig. 4b,c). This similarity is expected given that both classifiers are trained to decode time- and decision-related information: the neural state that evolves during the time leading up to the button press should carry anticipatory information about the decision to press the button. However, there were subtle (though not significant) differences between the signal distributions. For instance, sensorimotor signals may have contributed more to the outcome identity decoder than to the time decoder (Fig. 6b, left), possibly because a motor action occurred during the time bin used to train the outcome identity decoder. Meanwhile, MTL signals were important for both classifiers, (Fig. 6b, right), consistent with prior reports of time encoding in MTL regions (Tsao et al., 2018; Umbach et al., 2020).

**Figure 6.**
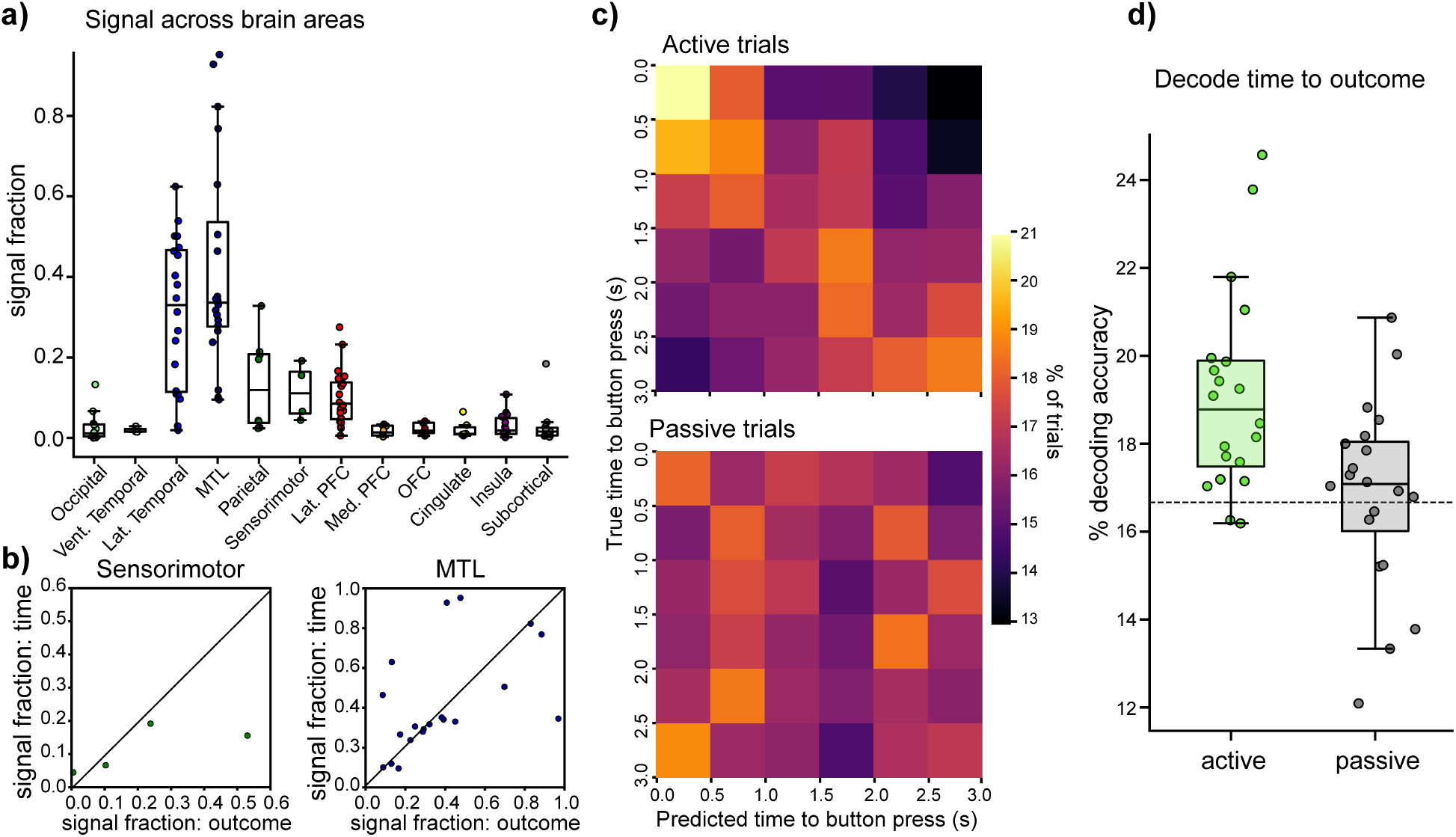
Linear decoding of absolute time to response. a) Spatial distribution of temporal signals (N = 20 sessions): signal carried by different brain regions (as in Fig. 4b). b) Contributions to the decoding of outcome identity (x-axis) compared to time-to-outcome (y-axis), for contacts in sensorimotor regions (left) and for medial temporal lobe contacts (right). c) Linear decoding of time-to-outcome (button press to stop inflation). N = 20 sessions. Confusion matrices illustrate time decoding for active and passive (control) trials. Colorbar shows the percent of trials in which the time bin on the y-axis is predicted to be the time bin on the x-axis. Time 0 is the last 0.5 second time bin before the subject’s response. d) Quantified decoding accuracy for active (green) and passive (grey) trials.

Interestingly, time-to-outcome was only linearly decodable on active trials (Fig. 6c, top; *p* = 0.00014), not passive (control) trials (bottom; *p* = 0.67) in which reward was guaranteed and no action was required to stop inflation. Decoding performance for active trials was significantly higher than for passive trials (Fig. 6d; paired *t*-test, *p* = 0.0022). Together, these results imply multiple facets of time processing in temporal and parietal regions, and that time coding of task-defined intervals only emerges when demarcated by a behaviorally relevant event.

### Neural signals are time-aligned to active decisions

Above, we demonstrated time decoding relative to an active decision on bank trials. We would not expect neural activity to predict outcome-aligned time on pop trials, when temporally structured neural signals anticipate an unexecuted decision planned for an unknown future timepoint. However, we were unable test this assumption, due to the following limitations of the SVM time decoding: 1) we only considered yellow balloon trials, to ensure >3 second inflation for most trials, and as a result, 2) we only attempted to decode time on bank trials, as there were not enough yellow pop trials.

Thus, to assess whether neural population activity contains temporally structured information sufficient to predict the timing of an imminent decision but not an unexpected pop event, we trained a nonlinear causal temporal convolutional network (TCN) to decode the time of trial response (Fig. 7a). This model allowed us to predict the moment of trial outcome based on cumulative data over the time course of the trial, for all three balloon colors. 4-fold cross validation showed that the TCN model can accurately predict the time of response specifically on bank trials (accuracy = .94 ± .012, MAE = 157 ± 52 ms) (Fig. 7b). The model’s prediction error was significantly higher for 3 individuals (Fig. S5a), and correlated with specifically longer trials that indicate distraction between cue and response (Fig. S5b).

**Figure 7.**
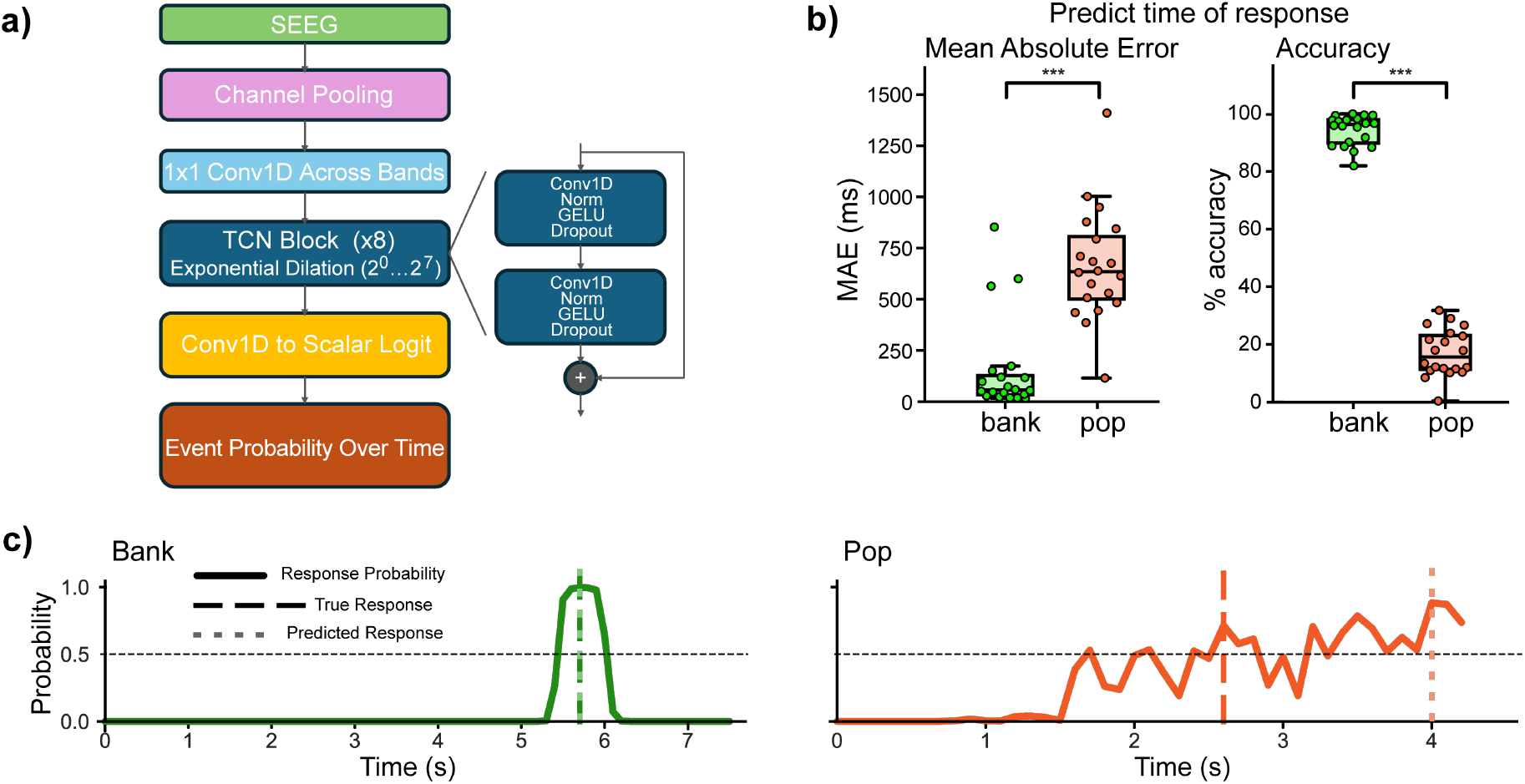
A temporal convolutional network predicts time of upcoming decision. a) Schematic of model architecture. b) Model performance (prediction accuracy) for predicting time of outcome for successfully executed decisions (bank trials) vs. unsuccessful pop trials. c) Probability of predicted response across the time course of an example bank trial (left) and an example pop trial (right).

As a control, we attempted to decode the time of pop events (which are stochastic and should not be represented by any brain signal). This control analysis would confirm that we are in fact decoding neural representations of temporal events rather than an artifact. Indeed, the TCN model could *not* predict the time of pop outcomes (accuracy = .17 ± .018, MAE = 669 ± 61 ms): we observed a significant difference in accuracy and MAE between TCN predictions of outcome time for bank vs. pop trials (*t*-test, p < 0.001) (Fig. 7b). Qualitatively, the TCN detected a gaussian of probability around the target event in active trials, but failed to identify a uniform probability around balloon pops (Fig. 7c). These results indicate that causal data accumulate in the LFP to allow prediction of meaningful choice behavior.

## Discussion

The present study found representations of time progression and temporal decision-related information in the human brain, by recording intracranial LFP signals in epilepsy patients performing a risky timing-dependent decision-making task. Beyond risk encoding, we identified a linearly decodable signal representing the presence or absence of a decision (bank and pop trials, respectively). Consistent with prior demonstrations that decision-related neural signals can precede motor actions (e.g., Libet et al., 1983; Fried et al., 2011), this signal extended backward in time at least 100 ms: a linear decoder trained at the time of the outcome could predict pop vs. bank outcome based on neural data 100 ms earlier.

Next, we sought to identify and characterize neural representations of time progression, which require temporally evolving structured trajectories in neural state space that track the passage of time. Note that the existence of such representations is not inevitable. Time would not be decodable if these trajectories were unstructured, inconsistent, or characterized by fixed-point dynamics (Cueva et al., 2020; Ahmed et al., 2020).

In line with rodent studies (Mello et al., 2015; Shimbo et al., 2021), we found temporally structured neural representations that scaled with trial duration: decoding after nonlinear dimensionality reduction revealed temporal information relative to inflation start and stop, regardless of the duration of this inflation period. To our knowledge, no prior studies have reported scalable neural representations of time in the human brain: although time-tuned cells have been identified in the human brain (Schonhaut et al., 2023; Umbach et al., 2020), these cells were observed to tile fixed, rather than scalable, durations.

In addition to relative timing, we uncovered representations of time passage aligned to an active decision. The progression of time leading up to the outcome/reward was only linearly decodable on active rather than passive trials. This observation agrees with previous findings, which report that time must be behaviorally relevant in order to be encoded (Rabinovich et al., 2022) and that the evolution of temporal dynamics depends on behavioral strategy and aligns with learned behavioral features (Bigus et al., 2024; Bowler et al., 2025). The present study provides direct evidence, through distributed human intracranial recording, that decision-aligned time coding is not simply a consequence of a temporally-evolving stimulus, but rather emerges when required for an active choice.

Importantly, these findings do not imply that participants relied on internal timekeeping alone. Since balloon size increased predictably over time, participants likely used visual feedback about the rate and extent of inflation to anchor their temporal predictions. We believe that this visual feedback does not supplant the role of timing, but rather facilitates it, since temporal prediction is critical for high accuracy in the task. Since the maximum balloon size is randomized across trials, responding based on only on balloon size would be an unsatisfactory strategy. Meanwhile, calculating instantaneous pop probability (based on a mental comparison between current balloon size and a internally-constructed goal template) is likely too slow to yield high accuracy. However, future experiments that decouple elapsed time from visual balloon inflation will be useful for distinguishing purely internal timing mechanisms from temporally structured sensory-feedback signals.

Inherent constraints of research in human patients also complicated our ability to pinpoint the precise functional-anatomical correlates of human time processing. The electrode placement was determined purely based on clinical need; therefore, the spatial coverage of our electrophysiological recordings was both limited and highly variable across participants. However, we were able to partially analyze the contributions of various brain regions to the encoding of temporal decision-making. Decision-related time information resided primarily in the temporal lobe—both medial and lateral regions. Time-keeping in the medial temporal lobe fits into conventional views about time encoding in this area. However, we speculate that time-related computations are actually more widespread, but that representations in other regions (e.g., sensory cortices) were hidden due to limited spatial coverage in our recordings.

Finally, we were able to recover high-resolution outcome-aligned time information, but only leading up to a planned decision. As expected, the timing of unanticipated pop events could not be predicted: on pop trials, any plans regarding an upcoming response would be interrupted by the pop, and neural signals tracking time would be aligned to some future, undetermined point.

Taken together, our results reveal that neuronal populations in the human brain track behaviorally-relevant time flow and encode temporally-structured information that predicts upcoming decisions made under risk.

## References

Ahmed, M. S., Priestley, J. B., Castro, A., Stefanini, F., Solis Canales, A. S., Balough, E. M., Lavoie, E., Mazzucato, L., Fusi, S., & Losonczy, A. (2020). Hippocampal network reorganization underlies the formation of a temporal association memory. Neuron, 107, 283–291.e6. 10.1016/j.neuron.2020.04.013.

Ashe, J., & Bushara, K. (2014). The olivo-cerebellar system as a neural clock. Advances in Experimental Medicine and Biology, 829, 155–165. 10.1007/978-1-4939-1782-2_9.

Bakhurin, K., Goudar, V., Shobe, J., Claar, L., Buono-mano, D., & Masmanidis, S. (2017). Differential encoding of time by prefrontal and striatal network dynamics. The Journal of Neuroscience, 37 (4), 854–870. 10.1523/JNEUROSCI.1789-16.2016.

Bigus, E. R., Lee, H.-W., Bowler, J. C., Shi, J., & Heys, J. G. (2024). Medial entorhinal cortex plays a specialized role in learning of flexible, context-dependent interval timing behavior. bioRxiv: The Preprint Server for Biology, 2023.01.18.524598. 10.1101/2023.01.18.524598.

Bowler, J. C., Azhar, D., Jensen, C. M., Lee, H.-W., & Heys, J. G. (2025). Structured experience shapes strategy learning and neural dynamics in the medial entorhinal cortex. bioRxiv, 2025.05.13.653873. 10.1101/2025.05.13.653873.

Clark, A. (2013). Whatever next? predictive brains, situated agents, and the future of cognitive science. Behavioral and Brain Sciences, 36(03), 181–204. 10.1017/S0140525X12000477.

Cueva, C. J., Saez, A., Marcos, E., Genovesio, A., Jazayeri, M., Romo, R., Salzman, C. D., Shadlen, M. N., & Fusi, S. (2020). Low-dimensional dynamics for working memory and time encoding. Proceedings of the National Academy of Sciences, 117, 23021–23032. 10.1073/pnas.1915984117.

Davis, T., Caston, R., Philip, B., Charlebois, C., Anderson, D., Weaver, K., Smith, E., & Rolston, J. (2021). Legui: A fast and accurate graphical user interface for automated detection and anatomical localization of intracranial electrodes. Frontiers in Neuroscience, 15. https://www.frontiersin.org/articles/10.3389/fnins.2021.769872

Fried, I., Mukamel, R., & Kreiman, G. (2011). Internally generated preactivation of single neurons in human medial frontal cortex predicts volition. Frontiers in Neuroscience, 69, 548–562 10.1016/j.neuron.2010.11.045

Gramfort, A., Luessi, M., Larson, E., Engemann, D., Strohmeier, D., Brodbeck, C., Goj, R., Jas, M., Brooks, T., Parkkonen, L., & Hämäläinen, M. (2013). Meg and eeg data analysis with mne-python. Frontiers in Neuroscience, 7. 10.3389/fnins.2013.00267.

Haufe, S., Meinecke, F., Görgen, K., Dähne, S., Haynes, J.-D., Blankertz, B., & Bießmann, F. (2014). On the interpretation of weight vectors of linear models in multivariate neuroimaging. NeuroImage, 87, 96–110. 10.1016/j.neuroimage.2013.10.067.

Larson, E., Gramfort, A., Engemann, D., Leppakangas, J., Brodbeck, C., Jas, M., Brooks, T., Sassenhagen, J., McCloy, D., Luessi, M., King, J.-R., Höchenberger, R., Brunner, C., Goj, R., Favelier, G., Vliet, M., Wronkiewicz, M., Appelhoff, S., Rockhill, A., & user27182. (2025). 10.5281/zenodo.17675410.

Lejuez, C., Read, J., Kahler, C., Richards, J., Ramsey, S., Stuart, G., Strong, D., & Brown, R. (2002). Evaluation of a behavioral measure of risk taking: The balloon analogue risk task (bart. Journal of Experimental Psychology. Applied, 8(2), 75–84. 10.1037//1076-898x.8.2.75.

Libet, B., Gleason, C. A., Wright, E. W., & Pearl, D. K. (1983). Time of conscious intention to act in relation to onset of cerebral activity (readiness-potential). The unconscious initiation of a freely voluntary act. Brain: A Journal of Neurology, 106 (Pt 3), 623–642. 10.1093/brain/106.3.623.

Mello, G., Soares, S., & Paton, J. (2015). A scalable population code for time in the striatum. Current Biology, 25(9), 1113–1122. 10.1016/j.cub.2015.02.036.

Pastalkova, E., Itskov, V., Amarasingham, A., & Buzsáki, G. (2008). Internally Generated Cell Assembly Sequences in the Rat Hippocampus [Publisher: American Association for the Advancement of Science]. Science, 321(5894), 1322–1327. 10.1126/science.1159775.

Pedregosa, F., Varoquaux, G., Gramfort, A., Michel, V., Thirion, B., Grisel, O., Blondel, M., Prettenhofer, P., Weiss, R., Dubourg, V., Vanderplas, J., Passos, A., Cournapeau, D., Brucher, M., Perrot, M., & Duchesnay, É. (2011). Scikit-learn: Machine learning in python. Journal of Machine Learning Research, 12(85), 2825–2830.

Rabinovich, R., Kato, D., & Bruno, R. (2022). Learning enhances encoding of time and temporal surprise in mouse primary sensory cortex. Nature Communications, 13(1), 1. 10.1038/s41467-022-33141-y.

Rao, H., Korczykowski, M., Pluta, J., Hoang, A., & Detre, J. (2008). Neural correlates of voluntary and involuntary risk taking in the human brain: An fmri study of the balloon analog risk task (bart. NeuroIm age, 42(2), 902–910. 10.1016/j.neuroimage.2008.05.046.

Salvioni, P., Murray, M. M., Kalmbach, L., & Bueti, D. (2013). How the Visual Brain Encodes and Keeps Track of Time. Journal of Neuroscience, 33(30), 12423–12429. JNEUROSCI.5146-12.2013.

Schneider, S., Lee, J., & Mathis, M. (2023). Learnable latent embeddings for joint behavioural and neural analysis. Nature, 617 (7960), 360–368. 10.1038/s41586-023-06031-6.

Schonhaut, D., Aghajan, Z., Kahana, M., & Fried, I. (2023). A neural code for time and space in the human brain. Cell Reports, 42(11), 113238. 10.1016/j.celrep.2023.113238.

Shimbo, A., Izawa, E.-I., & Fujisawa, S. (2021). Scalable representation of time in the hippocampus. Science Advances, 7 (6), 7013. 10.1126/sciadv.abd7013.

Toso, A., Reinartz, S., Pulecchi, F., & Diamond, M. (2021). Time coding in rat dorsolateral striatum. Neuron. 10.1016/j.neuron.2021.08.020.

Tsao, A., Sugar, J., Lu, L., Wang, C., Knierim, J., Moser, M.-B., & Moser, E. (2018). Integrating time from experience in the lateral entorhinal cortex. Nature, 561(7721), 57–62. 10.1038/s41586-018-0459-6.

Umbach, G., Kantak, P., Jacobs, J., Kahana, M., Pfeiffer, B., Sperling, M., & Lega, B. (2020). Time cells in the human hippocampus and entorhinal cortex support episodic memory. Proceedings of the National Academy of Sciences, 117 (45), 28463–28474. 10.1073/pnas.2013250117.

Wittmann, M., & Paulus, M. (2008). Decision making, impulsivity and time perception. Trends in Cognitive Sciences, 12(1), 7–12. 10.1016/j.tics.2007.10.004.

Young, D., Goodie, A., Hall, D., & Wu, E. (2012). Decision making under time pressure, modeled in a prospect theory framework. Organizational Behavior and Human Decision Processes, 118(2), 179–188. 10.1016/j.obhdp.2012.03.005.

