## Supplementary Materials for "Temporal Processing during Decision Making under Uncertainty"

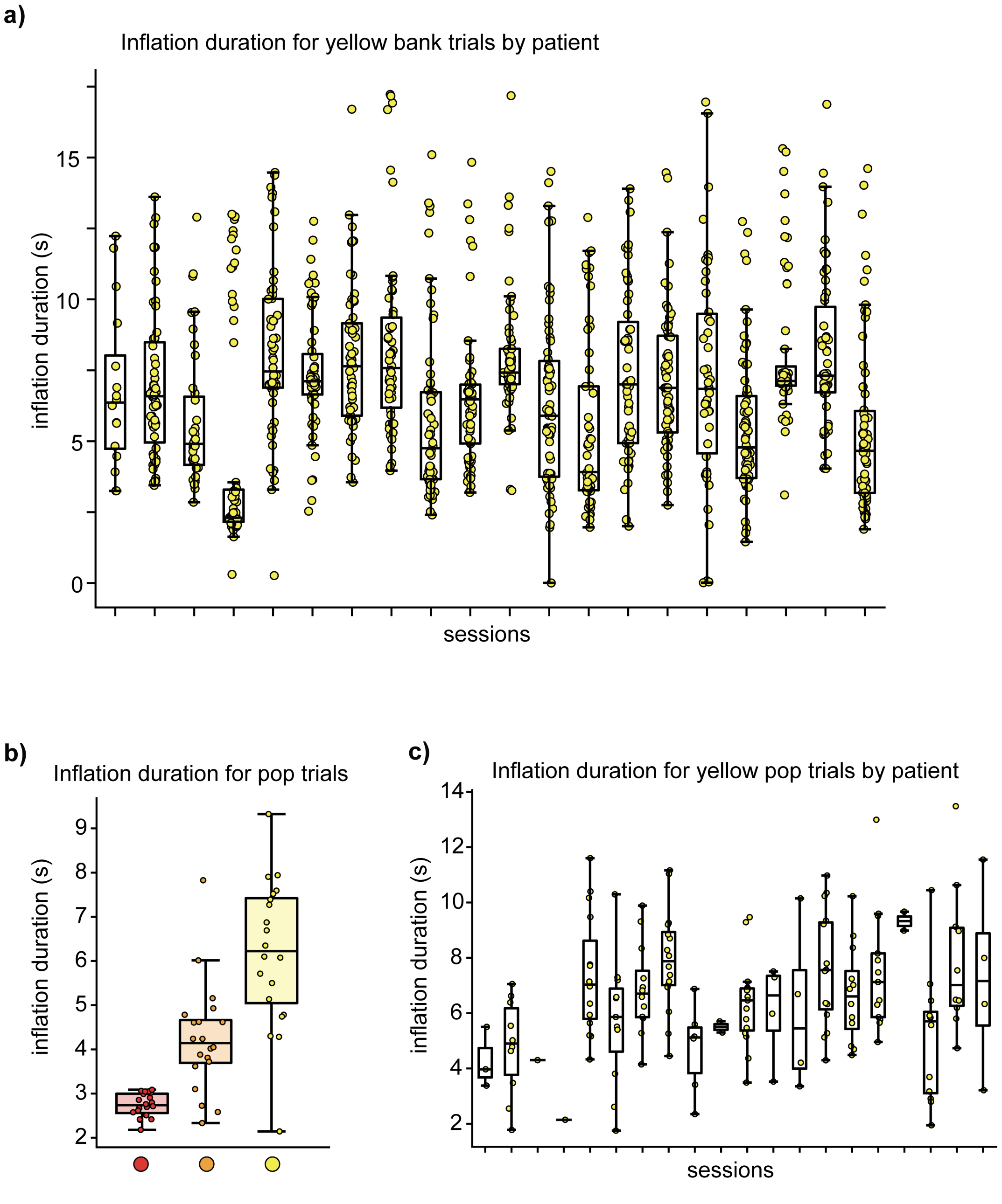


Figure S1: Characterization of behavioral responses. a) Duration of inflation for yellow-balloon bank trials, split by session. Each dot is a single trial. Note the variability in inflation times (i.e., time of button press) between trials within each participant's session. b) Duration of inflation for pop trials, for all three balloon colors. For each color, each dot is the average pop time for each patient. c) Same as (a), but for yellow-balloon pop trials.

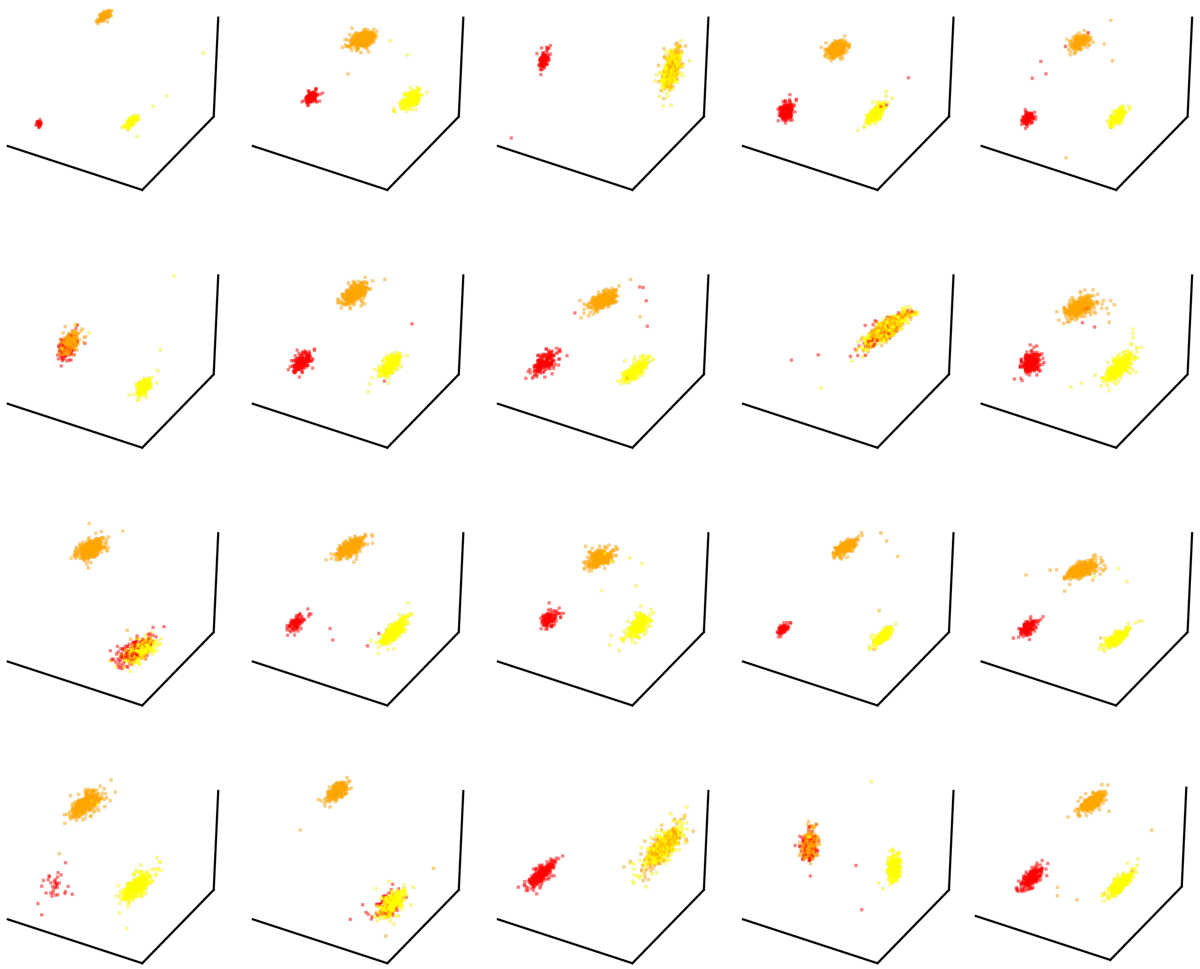

Figure S2: Single-session CEBRA-generated embeddings separating neural activiy related to different balloon colors during cue presentation. Each subplot illustrates the embedding for an individual subject or session.

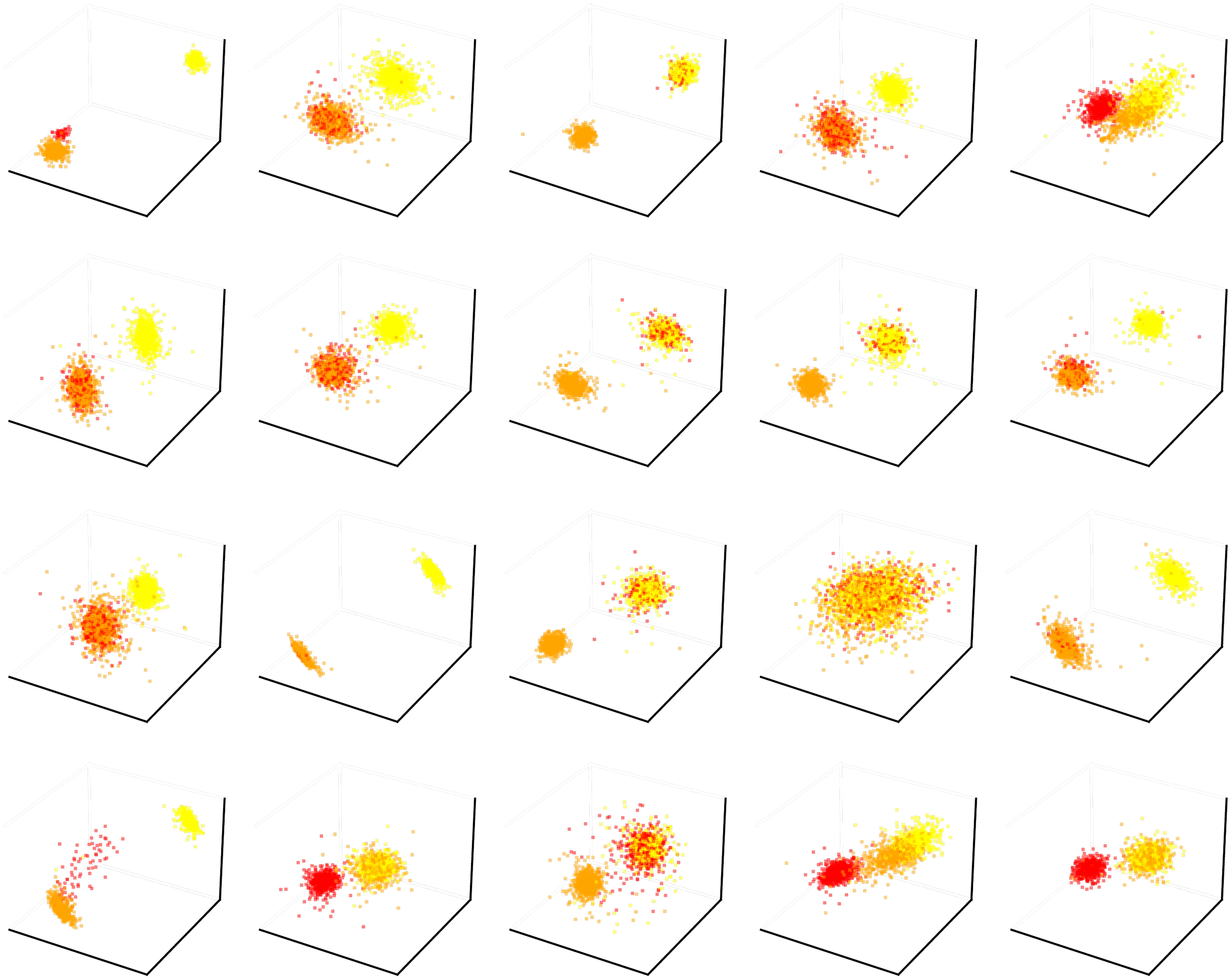

Figure S3: As in Fig. S2, but for the inflation period (3 seconds prior to response).

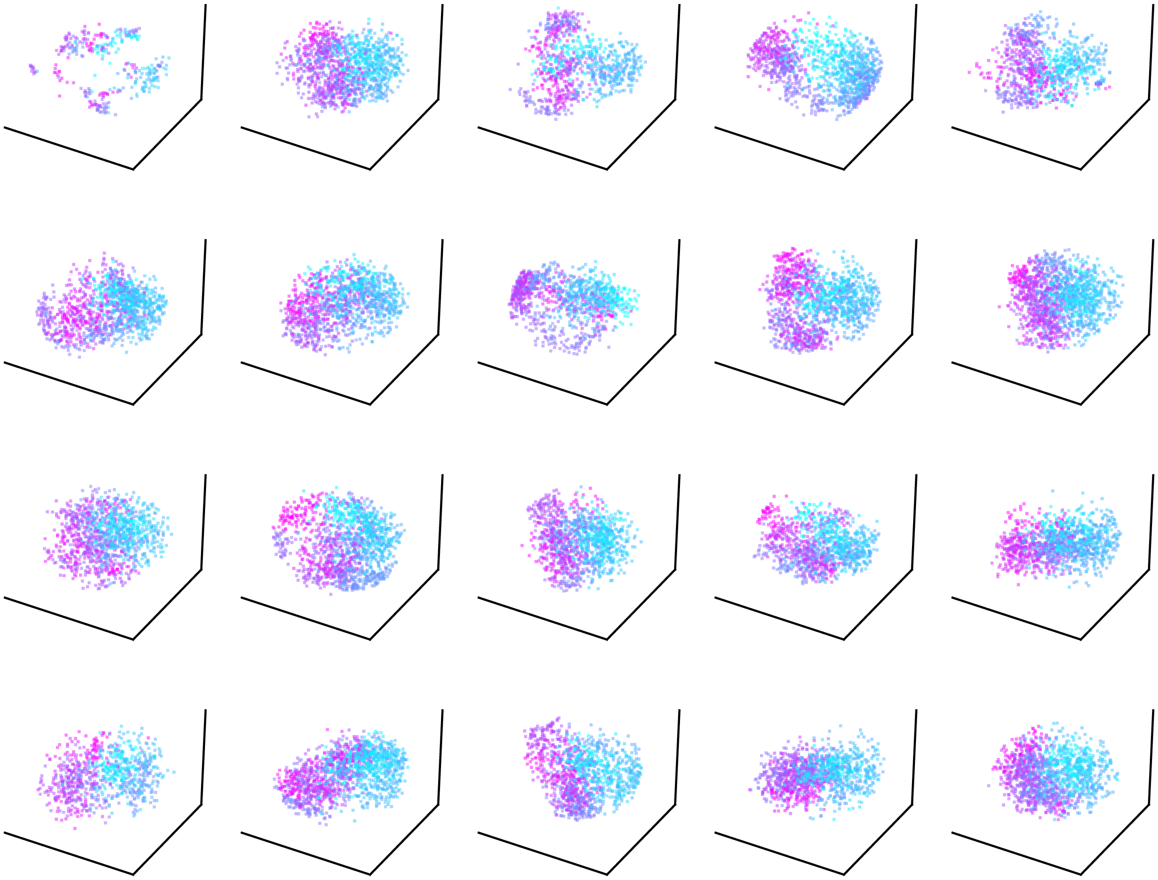

Figure S4: CEBRA-generated embeddings illustrating time-to-outcome for all 20 sessions. Each subplot is 1 session. Blue dots are timepoints closer to the button press that stops inflation

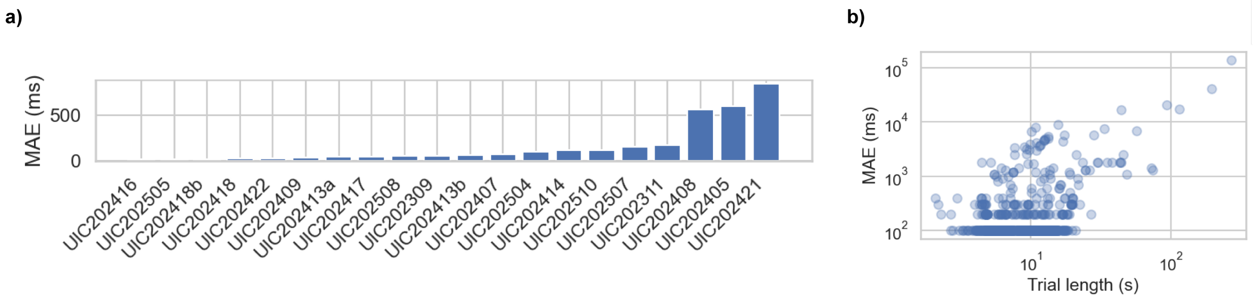


Figure S5: TCN prediction mean absolute error, by patient. a) TCN performance for each patient, sorted by MAE. b) MAE compared to total trial length (cue duration + inflation duration
